# Active segregation in binary mixtures under flow

**DOI:** 10.1101/2024.09.20.614032

**Authors:** Giacomo Di Dio, Remy Colin

## Abstract

Many bacterial habitats, from the gut to the soil, feature narrow channels where confined flow is a key constraint that might influence the spatial organization, and thus the functioning, of the complex, phenotypically heterogeneous communities these microbes form. Here, we investigate how a model heterogeneous bacterial community of motile and non-motile *Escherichia coli* organizes under confined Poiseuille flow. We discovered a new mode of active self-organization, where the motile bacteria induce the rapid sideways segregation of the non-motile cells to one side of the channel, eventually resulting in asymmetric biofilm formation. Our experiments and modeling elucidated the purely physical segregation mechanism: the rheotactic drift of the motile cells, which stems from shear acting on their chiral flagella, induces a conveyer-belt-like backflow advecting the non-motile cells. The latter then accumulate thanks to sedimentation countering flow incompressibility. This unexpected consequence of motility can affect the organization of complex bacteria communities colonizing environments under confined flow.

## Introduction

Natural bacterial communities usually form highly organized structures that are important for the functioning of the community, with strong impacts on health, ecology, and industry [1–3]. It is therefore essential to understand the principles that drive the organization in space and time of the diverse phenotypes, stemming from different species or differentiated clones, in these structures. Although biochemical factors are well known to be at play [2, 4], the physics of microbial communities is also increasingly realized to play important roles in shaping their spatial organization. This includes external physical constraints, chiefly shear flows experienced in many confined environments, e.g., the soil, aquatic sediments, or the gastrointestinal tract [5–9], but also sedimentation [10, 11], as well as physical interactions between bacteria, particularly when active motility drives the population far from equilibrium [12–14].

Most natural bacterial populations indeed feature motile bacteria, which are often mixed with non-motile bacteria in proportions that can vary widely between environments [13, 15, 16]. The most widespread motility is flagellar swimming, most studied in the model organism *Escherichia coli*, in which the micrometric bacteria propel themselves by rotating their helical flagella and randomly explore the environment by alternating between forward runs at 10 −100 *µ*m/s and short reorienting tumbles [17]. During swimming, low Reynolds number fluid flows are elicited by the flagellum and the cell body, which can be modeled at first order as an extensile (“pusher”) dipole of forces [18]. These flows induce long-range hydrodynamic interactions with other bacteria and the physical boundaries of the environment, which add to short-range interactions to affect single-cell and population behaviors [19]. The interactions with boundaries lead to swimmers accumulating at the surface [20–23], where they swim with a small inward tilt [24–28] and describe circular trajectories [29–32]. At population level, in addition to the well-known collective motion emerging in pure suspensions of swimmers at high density [33–35], these physical interactions have been recently found to also have the potential to shape more complex microbial communities, even at low cell densities [14, 36–39]. Notably, in mixed motile and non-motile bacterial communities, non-equilibrium fluctuating density patterns of the non-motile species emerge from the interplay between sedimentation and fluid flows induced by the circular swimmers located near surfaces [14].

Shear flow is known to strongly impact the physical single-cell behavior of both motile and non-motile bacteria, and should thus strongly modify the picture obtained in suspensions at rest. Under shear, rodshaped bacteria rotate following Jeffery’s orbits [40], which combines when swimming with the ability to cross streamlines to produce non-trivial behaviors, including cycloid trajectories [41, 42] and migration toward regions of high shear in bulk Poiseuille flow [43–47]. The shape of the flagellum, a left-handed helix during propulsion for *E. coli*, also leads to sideways rheotaxis: Shear induces a lift force on the chiral flagellum but not on the body, and the resulting rheotactic torque orients the swimmer towards the flow vorticity direction, thus generating a rheotactic drift perpendicular to the flow [48–50]. At surfaces, the interaction of the swimmer with the flow and the boundary further leads to an upstream reorientation (weathervane-like effect) that produces upstream motility at low to moderate flow rates, whereas bacteria are advected downstream if the flow is too high [51–53]. The tilt toward the surface is reduced [54], which is associated with reduced surface trapping [55]. Finally, chirality-induced rheotaxis still prevails for low to moderate flow rates [51, 56–59], but oscillations can emerge at very high shear under the combination of the former surface effects [54, 59]. How these single-cell effects play on the organization of a multispecies community remains however relatively underexplored. Most studies at the population level under flow have focused on how shear flow mechanically reshapes surface-attached bacterial biofilm colonies [7, 8], which may stretch, redistribute, and ripple [60, 61], notably in complex geometries [62–64], with high shear enhancing or disrupting biofilms depending on its dynamics [65–68]. In multispecies communities, the focus has primarily been on the reshaping of the landscape of chemical interactions by flow, which affects their composition, dynamics, and evolution [69–74].

Despite the importance of flow for heterogeneous microbial communities and the latter showing specific physics-driven self-organization behaviors even at rest, the physics of microbes under flow is thus almost exclusively studied at the level of individual swimmers in homogeneous populations. Here, we addressed how the physics of active systems affects the organization of heterogeneous communities under flow. We used a binary mixture of motile and non-motile *E. coli*, which we previously established as a tractable model system for heterogeneous bacterial communities [14]. We discovered that the mixture actively segregates in a microfluidic Poiseuille flow. Non-motile cells are advected to the “left” side of the channel (i.e. opposite the vorticity direction) where they accumulate, at a speed that depends on motile cell density and flow rate. We demonstrate experimentally and in simulations that this transport is caused by a backflow that results from the collective chirality-induced rheotactic drift of the surface-oriented motile cells. We also show that non-motile cell accumulation requires sedimentation, which counters the incompressibility of the conveyerbelt-like backflow, to take effect. Finally, we show that this rapid accumulation leads, as the population grows at long times under flow, to an asymmetric channel colonization through biofilm formation, demonstrating the relevance of this effect to long-term microbial community organization.

## Results

### Leftwards segregation of non-motile bacteria in binary mixtures under flow

To isolate the physical effects of shear flow on the spatial organization of heterogeneous bacterial communities, we employed as a model system a binary mixture of a motile and a non-motile strain of the bacterium *E. coli*. To minimize biological complexity, the strains derive from the same ancestor and differ only in the deletion of the flagellin gene *fliC* in the non-motile strain and their tagging with different fluorescent markers (mNeonGreen for non-motile, mCherry for motile). We mix the individually pre-grown strains in controlled proportions in a no-growth motility buffer. The mixture is subjected to Poiseuille flow in a microfluidic channel (height H = 56 *µm*, width W = 1 mm, total length L = 370 mm), where long straight sections are connected by short serpentine meanders [Fig. 1a].

**Fig. 1:**
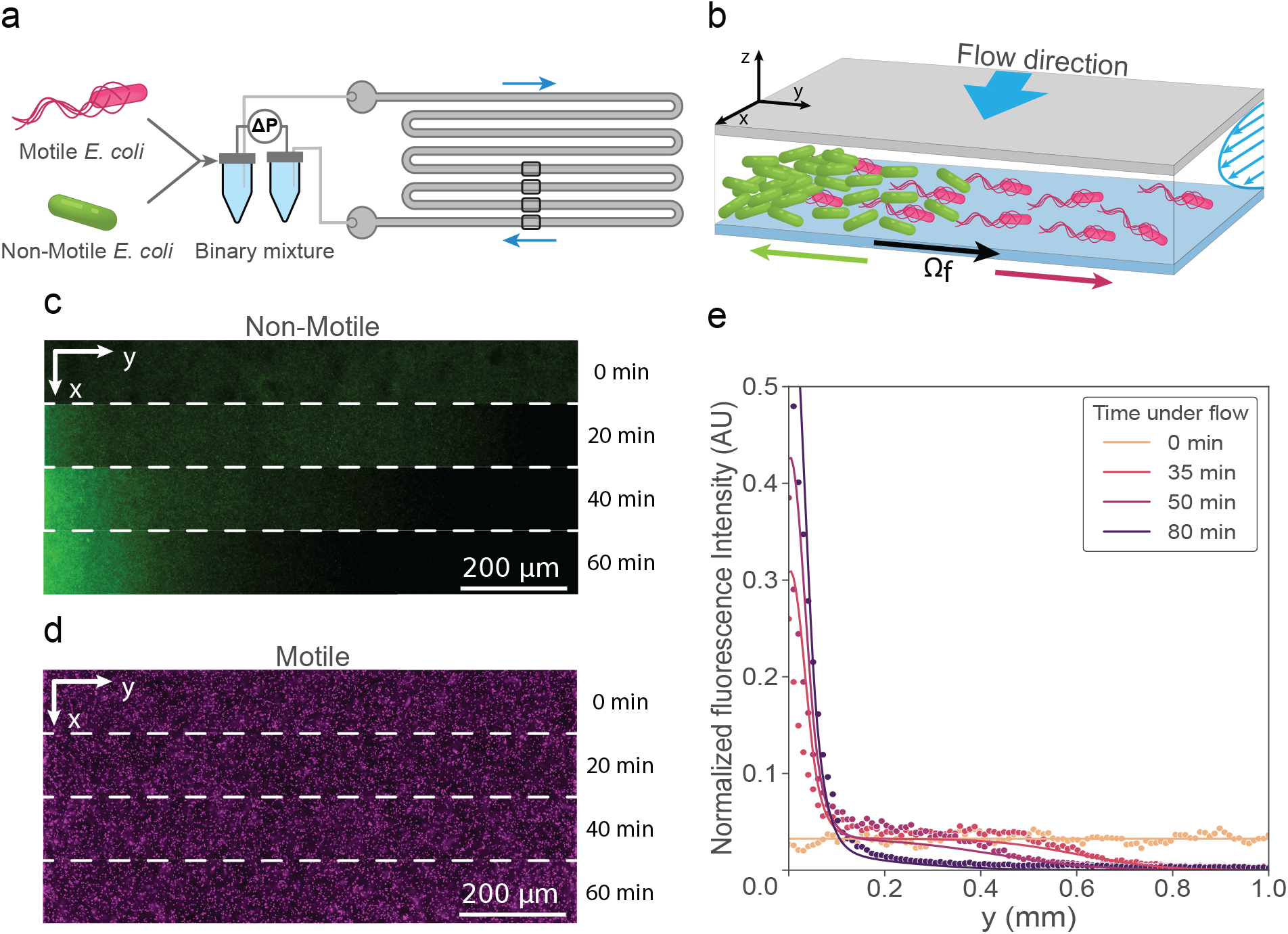
Active leftward segregation of the non-motile bacteria in binary mixtures under Poiseuille flow. **a** Sketch of the experimental set-up. The spatial organization of the mixture of motile and non-motile *E*.*coli* is observed under flow at the highlighted positions throughout the channel. **b** Schematic side view of the microfluidic channel. The ‘left’ side of the channel is defined as facing the flow direction x, i.e. opposite the direction y of the shear vorticity **Ω**_*f*_ at the bottom of the channel. **c,d** Example channel cross-sections of the non-motile (**c**) and motile (**d**) bacterial density, measured via fluorescence at indicated times and 10.5 *µ*m above the bottom surface. **e** Non-motile density profiles across the channel width at indicated times for a representative experiment (dots; *ϕ*_*M*_ = 0.17%, *ϕ*_*NM*_ = 0.17%, shear 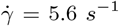) and the corresponding simulation of the one-dimensional advection-diffusion equation Eq. 2 (continuous lines; *v*_*y*_ = 0.19 *µ*m/s, *D* = 4.0 *µ*m^2^/s).

We follow the spatial organization of the mixture via time-lapse fluorescence microscopy over two hours close to the bottom (*z* = 10.5 *µ*m), where the non-motile cells sediment. We observed that the non-motile cells gradually re-distribute and accumulate on the “left” side of the channel when facing the upcoming flow, corresponding to the opposite of the flow vorticity direction at this height [Fig. 1b,c, Supplementary Fig. S1a]. In contrast, the distribution of motile bacteria remained homogeneous [Fig. 1d and Supplementary Fig. S2]. Segregation is absent in control experiments without motile cells, indicating that it is driven by their swimming activity. It also results from a local effect: The non-motile density profiles indeed evolve simultaneously and identically across the different straight sections of the chip [Supplementary Fig. S1b], the accumulation dynamics is independent of the number of meanders, and a similar segregation dynamics is observed for a smaller channel width (W=0.5 mm) [Supplementary Fig. S1c].

We quantified the evolution of the density profile *ρ*(*y*) of non-motile bacteria under flow, which shows simultaneous leftward accumulation and rightward depletion of non-motile cells [Fig. 1e]. At low non-motile volume fractions (*ϕ*_*NM*_ ≤ 0.17%), this dynamics is well-described by a one-dimensional advection-diffusion equation [Fig. 1e]:

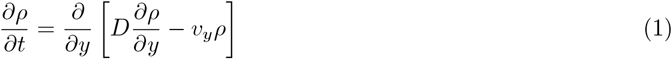

where the non-motile advection velocity *v*_*y*_ is constant and uniform throughout the bulk of the channel. It leads to an accumulation at the wall and is eventually balanced by the resulting diffusive counterflux (diffusion coefficient D) when reaching steady state. At higher concentrations of non-motile bacteria (*ϕ*_*NM*_ = 1.7%), although still qualitatively correct, this model does not quantify well the density profiles [Supplementary Fig. S1a], likely due to excluded volume effects at this very high non-motile density.

### The rheotactic motion of active swimmers drives the drift of passive non-motile cells

To understand the mechanism underlying this segregation, we analyzed how the non-motile advection velocity 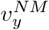 depends on possible control parameters. We measured 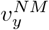 from the fluorescence density profiles via the center of mass of the distribution of non-motile cells along the width of the channel:

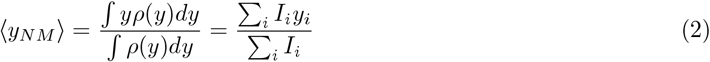

with *I*_*i*_ the mNeonGreen fluorescence intensity at pixel *i* with lateral coordinate *y*_*i*_, used as a proxy for the local non-motile density *ρ*(*y*). The center of mass *y*_*NM*_ starts at mid-channel, reflecting the initially homogeneous non-motile distribution, and drifts leftwards after flow actuation [Fig. 2a,b]. The drift velocity 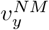 of the non-motile bacteria is the initial slope of ⟨*y*_*NM*_⟩(*t*), which we fitted with saturating exponential or linear functions depending on motile cell density [Fig. 2a]. Both smooth-swimming (Δ*cheY*) and run-and-tumbling (wild type) *E. coli* drive accumulation, the latter at a slightly lower rate [Fig. 2a,c]. This indicates that the rate at which swimming direction reorients is a control parameter of the system. Focusing on mixtures with smooth swimmers at a fixed flow rate, we found that the drift velocity first increases linearly with the motile volume fraction, before an inflection above *ϕ*_*M*_ = 0.17% [Fig. 2c]. Hence, the effect of the motile cells is mostly additive, except at higher density where motile cells appear to interact destructively and reduce their individual efficacy (see Discussion). The rate of left-side accumulation of non-motile bacteria was found to be independent of their concentration Φ_*NM*_ [Supplementary Fig. S3a]. Colloidal beads with a similar size to the bacteria also have a similar rate of accumulation [Supplementary Fig. S3b]. This indicates that the mechanism of segregation is purely physical and dependent on motile cell activity.

**Fig. 2:**
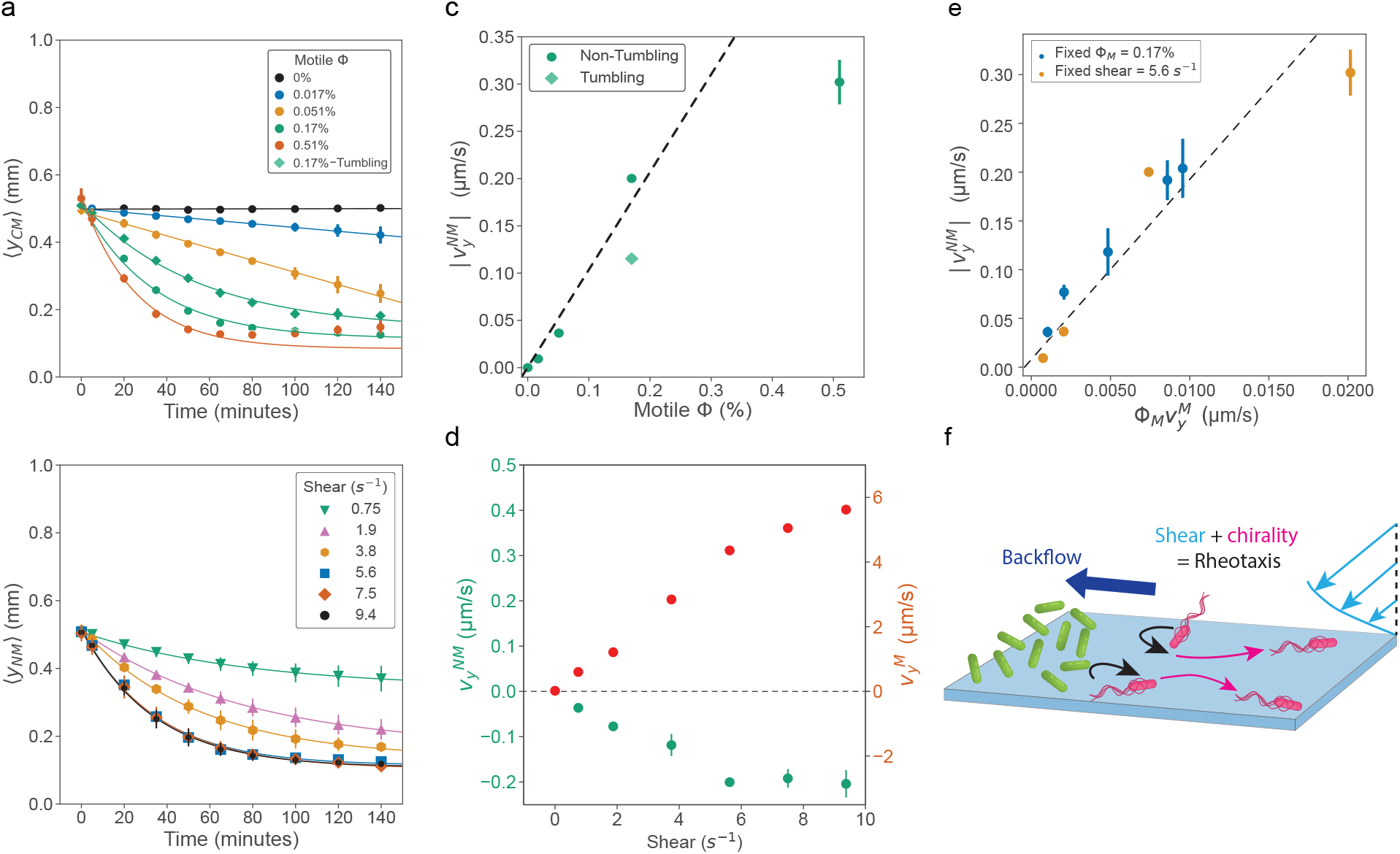
The rheotactic flux of motile cells induces the non-motile cell advection in the opposite direction. **a,b** Position of the center of mass of non-motile cells as a function of time for (**a**) different motile volume fractions (*ϕ*_*NM*_ = 1.7%, shear 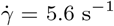), and (**b**) increasing values of applied shear (*ϕ*_*M*_ = 0.17%, *ϕ*_*NM*_ = 1.7%). Solid lines are linear (*ϕ*_*M*_ ≤ 0.051 % in **a**) or saturated exponential fits. **c** Drift velocity of the non-motile center of mass, obtained as the initial slope of the fitting curves of **a**. The dashed line represents a linear fit for non-tumbling data for *ϕ*_*M*_ ≤ 0.17% (slope = 0.96 *µ*m/s/%). **d** Drift velocities of non-motile cells 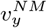, extracted from **b**, and the rheotactic velocity of motile cells 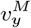, measured with particle tracking at 2.6 *µm* from the bottom surface (*ϕ*_*M*_ = 0.17%, *ϕ*_*NM*_ = 1.7%), as a function of shear. Positive y-direction is defined in Fig. 1b. **e** The relation between the drift of non-motile cells and the rheotactic flux of the motile cells 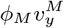. The dashed line represents a linear fit of all the points with a slope of 20.4. **a-e** Error bars represent the standard deviation (SD) over n=3 biological replicates. Each biological replicate is the average of four positions through the channel. **f** Schematics of the putative mechanism driving particle accumulation under shear flow.

The accumulation dynamics is also strongly affected by the flow rate experienced by the cells. We observed that the non-motile drift velocity increases linearly as a function of flow rate, quantified by the shear rate at the bottom surface 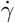, until it reaches a saturation above 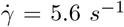 [Fig. 2b]. The accumulation dynamics was almost identical in experiments with a thicker microfluidic chip (H=80 µm) but the same shear rate at the surface 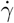 [Supplementary Fig. S4], showing that the latter is indeed the relevant control parameter. We tracked the motion of the motile cells close to the bottom surface (z = 2.6 *µ*m) and found that they performed a rheotactic drift in the vorticity direction (the ‘right’), as previously reported [48, 54]. The lateral drift velocities of the motile 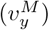 and the non-motile cells 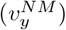 show a similar dependence as a function of shear rate [Fig. 2d], with 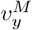 independent of volume fraction [Supplementary Fig. S5].

Bringing all our results together, we found that the drift of the non-motile cells 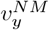 collapses as a single function of the lateral flux of motile cells 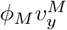, for all the volume fractions of motile cells and shear rates we probed [Fig. 2e]. This strongly suggests that the non-motile cells are advected by a backflow in reaction to the rheotactic drift of motile cells [Fig. 2f]. The motile cells, which tend to accumulate in equal proportions close to the top and bottom surfaces [Supplementary Fig. S6] [20, 21], generate low Reynolds number flow fields when swimming that are additive. The net swimmer flux due to chirality-induced rheotaxis in Poiseuille flow appears to generate a net backflow in the negative y-direction, which is hence proportional to the strength of the rheotaxis and the number of rheotactic cells [Fig. 2e]. The weaker non-motile drift induced by wild-type swimmers would then come from the reduced rheotactic drift of tumbling swimmers that randomize their swimming direction more rapidly [50]. However, two elements of this scenario remain to be elucidated: How is the backflow generated, and how does this incompressible backflow lead to non-motile cell accumulation?

### Sedimentation allows for non-motile bacteria accumulation by countering the conveyor-belt-like recirculation induced by motile cells

We first addressed the question of how the drifting non-motile cells accumulate. We previously showed that sedimentation is necessary for pattern formation in non-flowing binary mixtures [14]. We thus investigated the effect of sedimentation under flow. We prevented sedimentation of the bacteria by matching the volumetric mass of the suspending medium with that of the cells (1.1 g/cm^3^), supplementing it with the innocuous density-matching agent iodixanol. We observed that, under the same experimental conditions as previous experiments, the non-motile cell distribution stayed homogeneous in the absence of sedimentation [Fig. 3a].

**Fig. 3:**
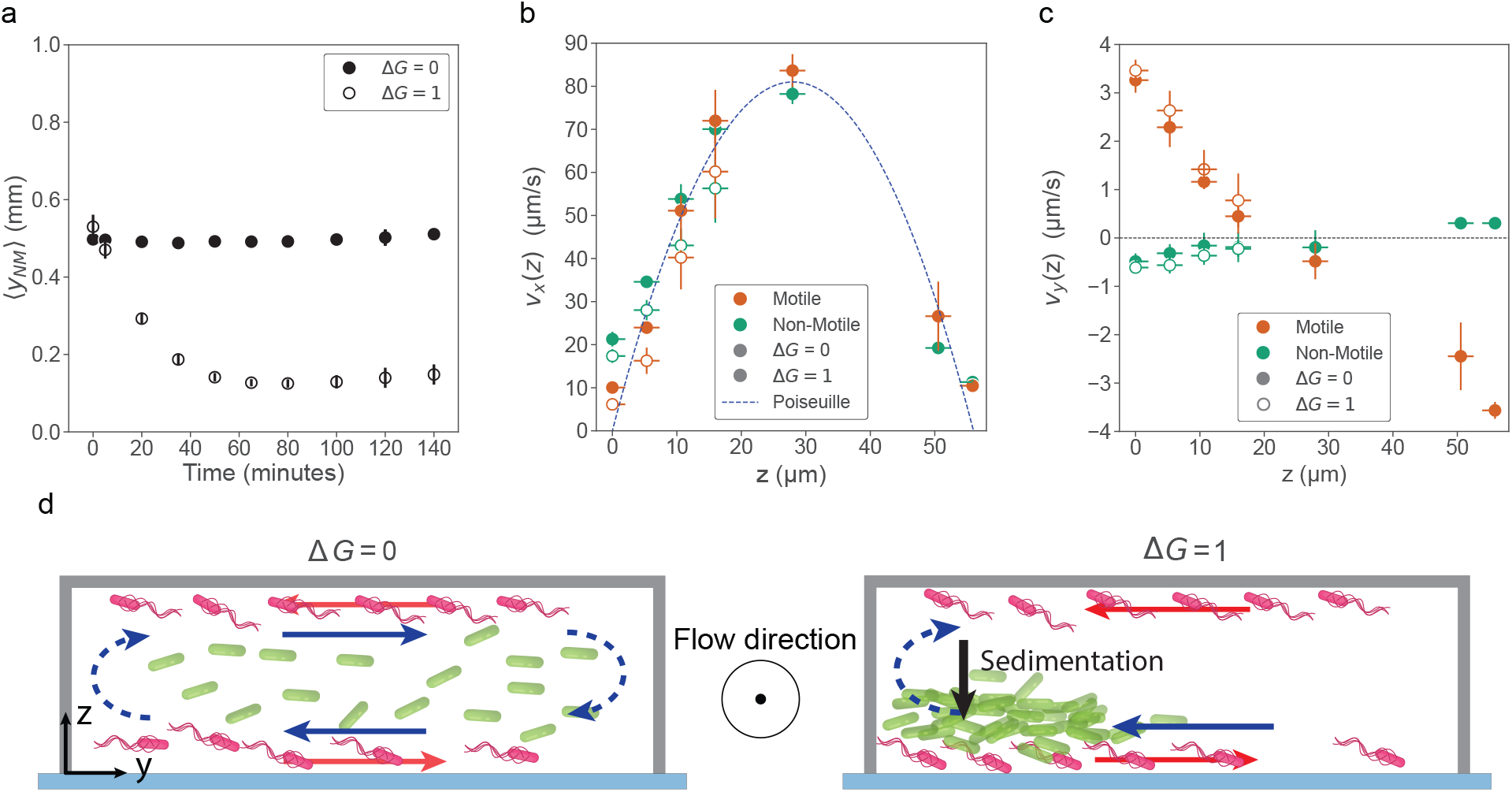
Sedimentation is necessary for the accumulation of non-motile bacteria that follow a recirculating backflow. **a** Position of the center of mass of non-motile bacteria as a function of time with (Δ*G* = 1) and without (Δ*G* = 0, density-matched medium) sedimentation (*ϕ*_*M*_ = 0.51%, *ϕ*_*NM*_ = 1.7%, 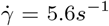). **b,c** Drift of motile and non-motile cells at increasing distances from the bottom surface, along (**b**) and perpendicular (**c**) to the direction of the main flow, with and without sedimentation, away from the side walls of the channel (*ϕ*_*M*_ = 0.51%, *ϕ*_*NM*_ = 1.7%, 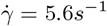). Vertical error bars represent the standard deviation (SD) over n=3 biological replicates, horizontal error bars represent the depth of field (4 *µ*m) of the fluorescence microscope. The drift is measured via *φ*DM, except for non-motile cells at z = 50.5 and 56 *µ*m where particle tracking was used to increase measurement precision (Methods). **d** Scheme of the accumulation mechanism. Perpendicular to the main flow, motile cells perform chirality-induced rheotaxis (red arrows), resulting in a recirculating, conveyor-belt-like, backflow that advects non-motile cells (blue arrows). Accumulation happens only in the presence of sedimentation (right).

Nonetheless, the non-motile cells kept being advected. We indeed measured the average drift of both motile and non-motile cells across the height of the channel using *φ*DM image velocimetry [75] and a high motile volume fraction (*ϕ*_*M*_ = 0.51%) to induce a large non-motile drift that can be accurately measured by this method. For both strains, there are no significant differences between the average drift velocities in sedimenting and non-sedimenting conditions, when they are measurable [Fig. 3b,c]. Indeed, non-motile drifts cannot be measured above *z* ≃ 20 *µ*m under sedimentation because few if any non-motile cells are visible in the field of view, while velocities are measurable throughout the channel in the non-sedimenting case. Along the main flow direction (*v*_*x*_(*z*)), both strains follow the Poiseuille flow profile [Fig. 3b], with minor deviations close to the surface as noted previously [76]. Perpendicular to the flow direction, the measurements captured the rheotactic drift of motile bacteria in the vorticity direction and the drift of non-motile cells in the opposite direction [Fig. 3c]. The rheotactic drift 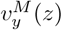 is an odd linear function of the distance to mid-height, being therefore proportional to the local shear rate as previously reported [48]. In particular, the motile rheotactic drift is oriented leftward at the top of the channel. The non-motile lateral drift 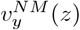 has an opposite sign to 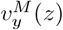, and increases as a function of z, also changing sign around mid-channel height, becoming rightward-oriented under the influence of the leftward-rheotactic motile cells at the channel top.

From these measurements in the density-matched suspending medium, we deduced that the backflow generated by the rheotactic motile cells takes the form of a recirculating conveyor belt in the y-z plane [Fig. 3d]. The fluid incompressibility indeed imposes that the recirculation flows upwards on the left side of the channel, and downwards on the right. Similarly to its role in the absence of flow [14], sedimentation then counteracts the upward flow, effectively compressing the non-motile cells and constraining them to accumulate on the left side. On the contrary, the motile cells are symmetrically distributed relative to mid-channel height, because swimming counteracts gravity [Supplementary Fig. S6], and they recirculate along the lateral walls, hence keeping a homogeneous distribution on the surfaces.

### Force dipole model explains the backflow generation by tilted rheotactic swimmers at surfaces

Since the extensile dipolar flow created by the motile bacteria should be fore-aft symmetric [18], it might be surprising at first that the rheotactic bias in the swimming direction induces a net backflow. However, most swimmers are located close to the surfaces [Supplementary Fig. S6], where swimming bacteria are known to be tilted a few degrees (*<* 10^*°*^) inwards [24–28, 76]. This tilt is also predicted to persist at our shear rates [54]. We note that short-range entrainment and repulsion by the swimmer as it moves can induce fluid transport, called Darwin drift, even for a swimmer parallel to the surface [77–80]. We used previous theoretical analyses of this drift for surface-aligned bacteria [78, 79] to estimate it to be about an order of magnitude lower than measured and therefore very unlikely to explain our observation [Supplementary Note 1]. We hence hypothesized that the tilt might be critical, by rendering the swimmer-induced fluid flow asymmetric, due to a larger effect of friction closer to the wall, and thus allowing the emergence of the backflow [76]. To test this hypothesis, we derived a model for the instantaneous flow generated by the bacteria.

We modeled the swimmers as a population of pusher force dipoles located close to the surfaces, with an inward tilt *θ* and a distribution of orientations in the x-y plane *φ* relative to the main flow [Fig. 4a]. We measured this distribution of *φ* from the instantaneous bacterial swimming directions in our experimental tracking data close to the bottom surface (Methods). It shows an increasing rightward (*φ >* 0) bias as shear increases, similar to previous bulk measurements [50], but also an upstream bias, which reflects the expected upstream rheotaxis [52] [Fig. 4b]. We place dipoles in equal numbers at the top and bottom surfaces, and we account for the change of vorticity direction at the top surface by a change of sign of *φ* for the swimmers there. The tilt angle *θ* is similarly symmetrized. We derived an analytical solution for the average Stokes flow generated between two no-slip walls by the swimmers, working in Fourier space [14, 81, 82] with open boundary conditions in the *x* and *y* directions, so that the model is valid far from the lateral walls of the channel.

**Fig. 4:**
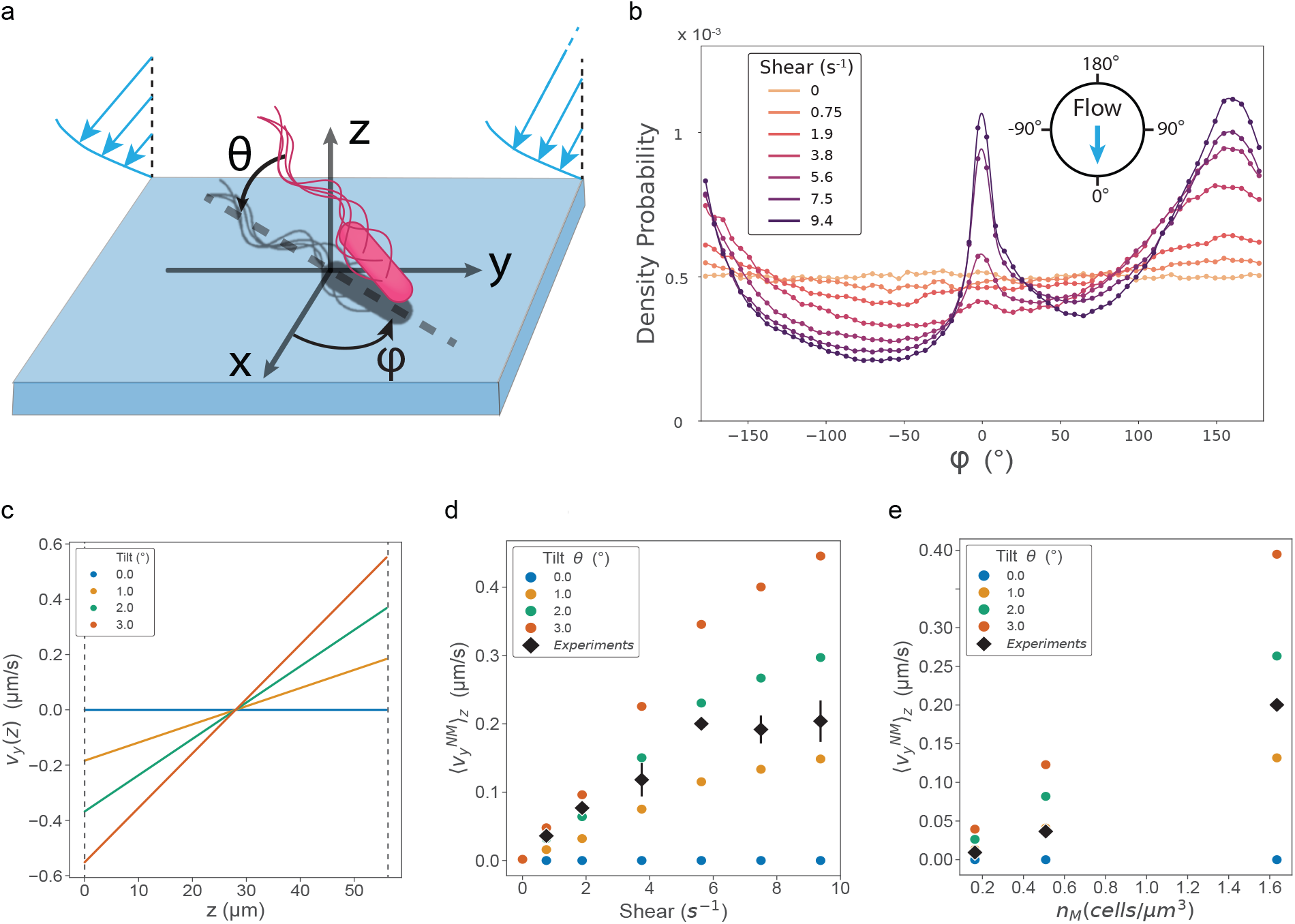
A model of rheotactic tilted force dipoles at surfaces explains the experimentally observed backflow generation. **a** System of reference for swimmers, with *φ* the x-y in-plane angle and *θ* the tilt towards the surface. **b** Distribution of the in-plane angle *φ* of smooth-swimming *E*.*coli* under indicated shear, measured with particle tracking in binary mixtures at 2.6 *µ*m from the bottom surface as the instantaneous velocity direction corrected from Poiseuille flow velocity. **c** Computed mean flow velocity in the y direction *v*_*y*_ (*z*) generated in the volume of the channel by a population of swimmers modeled as tilted force dipoles equally distributed across the upper and lower surfaces, for varying tilts *θ* toward the surface. Swimmers drift corresponds to a volume fraction *ϕ* =0.17 % and a shear 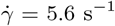. **d,e** Comparison between the predicted average drift of non-motile cells in the y direction, accounting for sedimentation (Eq. 26) and experimental results as a function of shear (**d**) and motile volume fraction (**e**). Error bars are standard deviation (SD) over 3 biological replicates for the experimental data.

The model predicts a swimmer-generated backflow in both the x and y directions (Methods). The sideward backflow 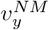 averaged over the x-y plane is a linear function of the z-position relative to mid-channel and directed toward the left (right) at the bottom (top) of the channel:

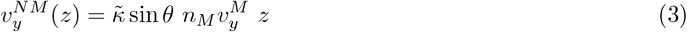

here, 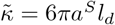 is the normalized dipole strength – with *a*^*S*^ the hydrodynamic radius of the cell body and *l*_*d*_ the dipole length, *n*_*M*_ = Φ_*M*_ */V*_*c*_ is the motile cell number density, *z* ∈ [−*h*, +*h*] with *h* half the channel height, and 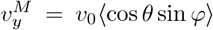 the chirality-induced rheotactic drift of the motile cells at the bottom surface. The backflow being non-zero thus requires swimmers to be tilted towards the surface (*θ* ≠ 0), which makes the dipolar flow asymmetric, and it increases with tilt, yielding realistic flow values for realistic tilts of a few degrees [Figs. 3c, 4c].

We next estimated from the predicted fluid backflow the non-motile drift in the presence of sedimentation, and its dependence on swimmer density and shear. The population-averaged non-motile drift is a weighted average of the fluid backflow, ⟨*v*_*y*_⟩_*z*_ = ∫ *p*(*z*)*v*_*y*_(*z*)*dz*. The vertical distribution of non-motile cells is given by Boltzmann, *p*(*z*) ∝exp(™(*z* + *h*)*/L*_*sed*_), with the sedimentation length of non-motile bacteria *L*_*sed*_ = 8 *µm*, which we measured previously in the presence of motile cells [14]. Integrating Eq. 3 over the Bolzmann distribution yields:

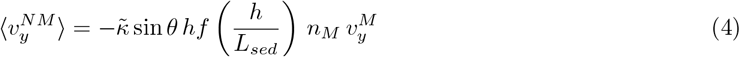

The function *f* (*h/L*_*sed*_) (Eq. 29) increases from 0 to 1 as *h/L*_*sed*_ increases from 0 (density match case) to infinity. The predicted drift ⟨*v*_*y*_⟩_*z*_ is thus non-zero when *p*(*z*) is skewed to the bottom, and it accurately reflects the experimental dependence of the non-motile drift both on shear rate [Fig. 4d] and on motile volume fraction for Φ_*M*_ ≤ 1.7% [Fig. 4e]. The expression also predicts the observed dependence in the bacterial flux 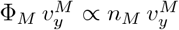 [Fig. 2c]. The model matches the experimental values, given our estimated value of 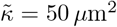, for a realistic tilt of *θ* ≃ 2^*°*^ [25–27, 54]. Note that the matching tilt decreases when the shear rate increases [Fig. 4d], in line with theoretical expectations [54].

The model also predicts a backflow in the x-direction in response to the upstream swimming of the motile bacteria (Eq. 24), 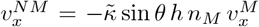, which is independent of sedimentation. This backflow should accelerate the downstream advection of the non-motile cells. However, combining Eqs. 24 and 25, we can estimate this predicted backflow as

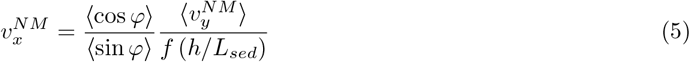

Given the measured distributions of *φ* [Fig. 4b], this yields a maximal 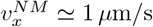 for the highest shear and volume fractions. This is well below the corresponding advection speeds by the main Poiseuille flow even fairly close to the surface [Fig. 3b]. It is also lower than our measurement error in this direction. Because motile bacteria swim upstream on all surfaces, we expect a segregation of motile and non-motile bacteria up and down the flow that should primarily result from this upstream motility, of order 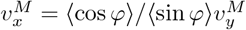 and which maximizes around 6 *µ*m/s for the same conditions, while the contribution of the backflow 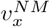 is secondary. However, we did not detect it in our system since we continuously inject bacteria into the channel.

Although fairly simple, our model of the underlying physics thus captures well the main features of the system and notably highlights the importance of the tilt of the motile cells at the surface.

### Rheotactic-induced accumulation reshuffles biofilm formation

Finally, we explored the implications of rheotaxis-induced accumulation for microbial communities at long time. For this, we used a binary mixture of a wild-type and a non-motile *E. coli* strain that both are capable of aggregating via *curli* fibers and antigen 43 (Ag43) expression. As the mixture grows over 48 hours at 30^*°*^ C in the microfluidic device under shear flow (*γ*? = 5.6 s^−1^), we observed that the non-motile fraction accumulates at the left side of the chip, advected by the rheotactic backflow, and forms an asymmetric biofilm at the edge of the channel [Fig. 5a]. This asymmetric biofilm formation is absent in controls with homogeneous populations of non-motile bacteria, which form biofilm uniformally on the surface of the channel [Fig. 5b]. The composition and development of the biofilm are influenced by the ability of motile bacteria to aggregate through the expression of Ag43. When motile cells can aggregate, they gradually integrate within the asymmetric biofilm produced by non-motile cells, with most motile cells eventually localizing within the biofilm [Supplementary Fig. S7a]. In contrast, when the motile bacteria cannot express Ag43, they remain distributed as single cells throughout the channel even after an extended period of time, as expected in our growth conditions [83, 84]. The biofilm is then almost exclusively composed of non-motile cells, it is generally smaller and less regular in thickness, consistent with the known structural role of flagella in forming robust biofilms in *E. coli* [85], but it remains strongly asymmetric [Supplementary Fig. S7b]. In both cases, our observations thus show that the interplay between motility and flow can indeed significantly affect over long time scales the patterns of biofilm formation in heterogeneous microbial communities.

**Fig. 5:**
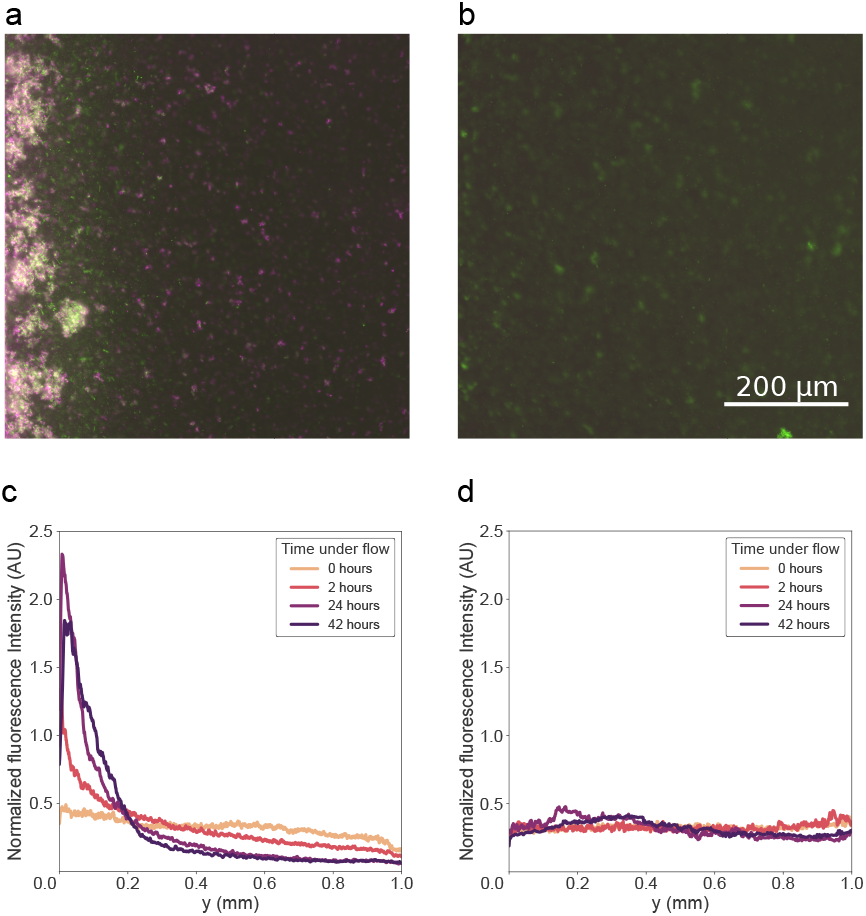
Rheotaxis-induced accumulation leads to asymmetric biofilm formation in aggregating strains. **a** Biofilm formation on the left side of the channel in a mixture of wild-type motile and non-motile *E. coli* (initial volume fractions: *ϕ*_*M*_ = 0.17%, *ϕ*_*NM*_ = 0.051%). The mixture is imaged after 40 hours under a continuous shear of 5.6 *s*^−1^. **b** Homogeneous biofilm formation in a control with only non-motile bacteria. **c,d** Evolution of the density profile of non-motile cells in the channel for the aggregating mixture (**c**) and the control (**d**). In both cases, the profile is calculated as the median over two biological replicates of the fluorescence signal at the 4 positions of the chip highlighted in Fig. 1.

## Discussion

Fluid flows are a ubiquitous environmental constraint that strongly affects many facets of the behavior of complex microbial communities, particularly in confined and tortuous environments like the animal gut, plant vessels, or the soil [6, 7, 9]. In particular, the consequences of flow affecting the physics of individual bacterial swimming [51–54, 56–59, 59] on the structuration of complex, phenotypically, and/or genetically heterogeneous bacterial populations are still being unraveled. Here, we discovered a new type of population structuration under confined flow, where non-motile bacteria in mixed communities with motile ones segregate at the bottom “left” side of microchannels. Combining experiments and modeling, we uncovered the purely physical mechanism driving segregation. The motile cells, which drift rheotactically in the flow vorticity direction (“rightwards” at the bottom), induce a backflow because they swim with a tilt at the surface that produces asymmetric dipolar flows. Non-motile cell advection by this backflow combines with sedimentation, which breaks vertical symmetry and counters the fluid incompressibility, to allow the passive cells to cross streamlines and accumulate at the left side of the channel. Lastly, this relatively fast segregation mechanism cements at long time in the asymmetric formation of dual biofilms. Hence, this active mechanism drives the spatial organization of heterogeneous bacterial communities at multiple time scales and over a wide range of physiological densities and relevant shear values, showing its relevance for the many natural microbial communities and biofilms that experience flow.

Although we work at higher densities and over a wider range, our measurements of motile *E. coli* rheotaxis are in good agreement with previous works that were limited to very diluted regimes [48, 50, 51, 56], in particular for the dependence of the lateral drift of motile cells on local shear [Figs. 2d, 3c]. Furthermore, the distribution of orientations of the motile bacteria under shear, which we measured here at the surface, is relatively similar to the one previously measured in bulk [50]. This distribution is also consistent with previous simulations of surface rheotaxis [54], except for the peak of probability of downstream motion that we observe at higher shear [Fig. 4b], due to downstream swimmers situated further above the surface that Mathijssen *et al* do not consider [54]. For a fixed chiral shape of the flagellum, the strength of rheotaxis depends on the Peclet number 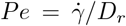 [50], explaining why tumbling, which increases the effective rotational diffusion coefficient *D*_*r*_, reduces the elicited non-motile drift [Fig. 2a,c]. Since the rheotactic velocity of motile cells is independent of their volume fraction [Supplementary Fig. S5], the saturation of 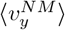 at high motile cell density Φ_*M*_ [Fig. 2] likely comes from the decrease in the fraction of motile cells located at the surface in this case [14].

The presence of backflow in the channel and the advection of non-motile bacteria against the motile cell motion contrasts with previous models and experiments in which passive tracers are entrained towards the mean direction of motion of active droplets [80]. The segregation mechanism thus provides a different mode of active transport of passive cargoes [86], that notably allows isolating passive from active particles [49, 87]. This difference comes from contrasting mechanisms, with bacteria inducing an asymmetric dipolar flow field by swimming at the surface with a tilt, which results from short-range interactions with the surface [24–28], whereas a volumetric effect, the Darwin drift, is dominant in the other system [77, 80]. This latter entrainment effect has been studied for bacteria models [78, 79]. Its sign and magnitude depend on the magnitude of the dipolar and quadrupolar terms in the expansion of the flow fields induced by the bacterium [78, 79]. Although it could be in the correct direction, we estimate the predicted drift induced by a swimmer parallel to the surface to be an order of magnitude lower than our measurement [Supplementary Note 1]. The tilt near the surface, which was absent in these previous models [78, 79], is therefore necessary to quantitatively explain our observations. Our model also predicts an enhanced downstream advection of the non-motile cells in response to the upstream swimming of the motile cells (Eq. 24). However, its magnitude is very small, consistent with previous simulations [76], suggesting that the differences in *v*_*x*_ between motile and non-motile cells close to the surface (Fig. 3b) primarily come from upstream motility and the finite depth of field of the microscope, rather than downstream entrainment. Our model took a simple approach, assuming that swimmers are force dipoles that are all located at the surface, neglecting steric interactions and higher-order terms in the flow field expansion. We neglected bacteria in the bulk, although shear could confer an average tilt to them, which would then contribute to the rheotactic backflow [43]. Assuming a constant tilt (∼ 2^*°*^) fits the experiments relatively well, although our results are also consistent with the expected reduced tilt at higher shear [44], and the fitted tilt values are close to those expected from simulations [54]. This indicates that despite its simplicity, the model captures the main physics of the system.

We believe our findings are relevant to a wide range of microbial communities in many confined environments, in particular the soil, plant roots, or aqueous sediments. Indeed, the accumulation mechanism is effective in a wide range of shear values and physiological volume fractions of motile cells. We observed it in channels of a few tens of microns height, which corresponds to the typical mode of the very broad distribution of mesopore sizes in the soil [88, 89] and to typical plant vessel sizes [90], and we found it to be independent of the channel width. Since the constraint of gravity is central to the mechanism, we expect segregation to occur in the relatively horizontal portions of natural porous networks (eg., the lateral roots of plants). Moreover, since the rheotactic mechanism is conserved among many motile bacterial species [48], so would the rheotaxis-induced accumulation and biofilm asymmetrization. Many (but not all) known bacterial flagella are left-handed, suggesting that a net segregation of non-motile cells could occur even in highly diverse bacterial communities. The asymmetric biofilm formed by the segregated non-motile cells can incorporate motile cells, only if they express Ag43 as observed in static cultures [83, 84], to form a robust mixed biofilm. In this case, the motile cells thus constructed via pure physics a habitat in the channel that required the non-motile partner, because motile biofilm formation was otherwise weak. This effect might benefit the many species that split between motile and non-motile subpopulations upon entry in biofilm-prone stationary phase [13]. Beyond facilitating biofilm formation due to the increased local cell density, the accumulation on the side of the channel, where shear is lower, can also protect the community from flushing by high shear [65, 66]. In summary, the results presented here demonstrate how shear may specifically affect the spatial organization of heterogeneous bacterial communities over time.

## Methods

### Strains

All strains are derivatives of *Escherichia coli* wild-type strain W3110 (RpoS+) and are listed in Supplementary Table S1. Apart for biofilm growth, strains lack the gene *flu* that encodes for the protein Ag43 mediating cell-cell adhesion to prevent confounding effects of cell aggregation when analysing segregation dynamics [91]. If not otherwise stated, the motile strain is a Δ*cheY* non-tumbling, smooth swimming, mutant. The nonmotile strain is a Δ*fliC* mutant that lacks flagellar filaments. The fluorescent markers are expressed from a plasmid carrying the fluorescent protein gene under the control of the Isopropyl-*β*-d-1-thiogalactopyranoside (IPTG)-inducible Tac promoter and an ampicillin (Amp) resistance marker. The non-motile strain is tagged with mNeonGreen and the motile strain with mCherry [14]. We have extensively characterized these strains in previous works [14, 35, 91]. The wild-type motile strain swims at about *v*_0_ = 23 *µ*m/s. The Δ*cheY* mutant swims faster, at *v*_0_ = 30 *µ*m/s on average, as measured from particle tracking in our conditions. The cell bodies have a length of 2.5 *µ*m on average, a width of 0.8 *µ*m in our growth conditions. The volume fractions are computed from the optical density at 600 nm (OD_600_) measured with a previously calibrated photometer as Φ = 1.7 10^−3^× OD_600_ [35].

### Microfluidics and microfabrication

The microfluidic chip consists of a serpentine channel with rectangular cross-section (height *H* = 56 *µ*m and width *W* = 1 mm if not otherwise stated), made of long straight parts (length L = 40 *mm*) connected by curved meanders for a total channel length of several hundred millimeters. Molds were fabricated using standard soft photolithography techniques. The SU82050 photoresist (Microchem^*T M*^) was spin-coated on a silicon wafer, covered with a negative mask produced using AutoCAD and printed by JD Photo Data (UK), exposed to UV light, baked, and developed according to manufacturer instructions, and silanized. Chips are made of poly-di-methylsiloxane (PDMS, Sylgard 184, Dow) in a 1:10 crosslinker-to-base ratio, which was poured on the cast, degassed, baked overnight at 65°C, peeled off, cut to shape, hole-punched and covalently bound on isopropanol-rinsed microscopy glass slides after oxygen plasma treatment.

PDMS to glass covalent bounds were allowed to form for 25 minutes at room temperature. The device was then primed with the binary mixture solution to prevent bubble formation, before connecting it to the pressure pump. For biofilm experiments, the mixture was instead injected 4 hours after the bonding to increase surface hydrophobicity and favor adhesion of the bacteria to the walls of the chip. The flow in the channel is actuated using an external pressure pump (Elveflow OB1 MK4 microfluidic flow controller, Elvesys), by applying a constant pressure difference in the range of 0.2 ™ 0.9 mbar between the inlet and outlet of the chip. We verified the shear rate at the surfaces by measuring the maximum advection speed *v*_*x*,max_ at the mid-height of the channel using the non-motile cells as tracers.

### Segregation dynamics experiments

Motile and non-motile cells are grown separately overnight (ON) in tryptone broth (TB) medium (1% tryptone + 0.5 %*wt* NaCl) and 0.1 g/L Amp at 37°C. The ON culture is diluted 1*/*100 in 10 mL fresh TB supplemented with 0.1 g/L Amp and 100 *µ*M IPTG. The strains are then grown separately in a shaking incubator at 270 rpm and 34°C for 4 hours. The bacteria are washed and re-suspended in motility buffer (MB - 3.914 g/L NaCl, 10 mM KPO_4_ buffer at pH 7, 0.1 mM Ethylenediaminetetraacetate) supplemented with 0.01% Tween80 to prevent adhesion to surfaces and 1% glucose to ensure sustained motility throughout the experiment. In experiments with density-matching media, the cells are instead re-suspended in a solution of motility buffer supplemented with 20% iodixanol (*ρ* = 1.11 g/ml). The *OD*_600_ of the resuspended cells is measured and the desired concentrations of motile and non-motile cells are obtained by further dilutions and mixing in appropriate amount. The resulting binary mixture is continously injected into the microfluidic channel throughout the experiment. For experiments with beads, the mixture is prepared using the same protocol, substituting non-motile cells with FluoGreen-tagged PMMA particles of diameter 1.33 *µ*m ± 0.1 *µ*m (Microchem GmbH), and the mixture is initially injected into the channel but flow is the generated by injecting MB to save material.

### Biofilm experiments

The bacteria used for the biofilm experiments are *E. coli* wild type (RC 144) or Ag43 knockout (RC101 - Supplementary Fig. S7), both tagged with mCherry, for the motile strain and *E. coli* Δ*fliC* tagged with mNeonGreen (SEB 21). The strains are grown separately overnight in TB with 0.1 g/L Amp at 37^*°*^ C. Non-motile cells are then diluted 1*/*50 in M9 medium (Na_2_HPO_4_ 48 mM + KH_2_PO_4_ 22 mM + NaCl 8.4 mM + NH_4_ Cl 18.6 mM + MgSO_4_ 2 mM + CaCl_2_ 1 mM) supplemented with 0.4% succinate and 0.2% casaminioacids, 0.1 g/L Amp and 100 *µ*M IPTG. The motile strain is prepared in TB as described in the previous section. Cells are grown at T = 30°C, where they can form robust biofilms [92], in a shaking incubator at 270 rpm. The motile strain is grown for 5 hours, and the non-motile strain for 8 hours to ensure that bacteria are in the late exponential/early stationary phase where *curli* expression is high. The bacteria are then washed and re-suspended in the filtered spent M9 medium in which the non-motile strain was grown, since it promotes biofilm formation, in contrast to fresh medium that induces biofilm-repressing exponential growth. The mixture is injected into the microfluidic chip and left to rest for one hour, before flowing filtered spent M9 medium into the channel. The evolution of the mixture is observed at several positions through the channel for a total duration of 42 hours under a constant applied pressure, yielding a measured shear of 5.6 *s*^−1^ at the beginning of the experiment, and at T = 30°C.

### Images and video acquisition

The images and videos are acquired using a Nikon TI-E inverted fluorescence wide-field microscope with a 20× objective (NA 0.75), mCherry (excitation filter 572/25, emission 645/90) and GFP (excitation filter 470/40, emission 525/50) filter cubes, and an Andor Zyla sCMOS Camera (1 px = 0.35 *µ*m), piloted via the Nikon NIS software. The depth of field of the objective is around 4 *µ*m. Focus is maintained using the Nikon perfect focus system. Since the objective is dry, we correct the heights measured with the micrometric skrew of the microscope by the refractive index of water (n=1.33). For the measurements of bacterial drift, videos of 5 seconds at 100 fps are captured at different distances from the bottom surface away from the lateral walls. For the analysis of swimming trajectories with particle tracking, videos are captured at 50 fps for 40 seconds with a 2×2 pixel binning at 2.6 *µm* from the bottom surface.

### Density profile measurement and analysis

Images are captured at 6 different points (respectively at 25, 130, 210, 255, 300, and 345 mm from the inlet) halfway through the straight portion of the channel, at 10.5 *µ*m from the bottom surface. Images are recorded before (0 min) and every 5 minutes after flow actuation. Since the image size for 20x magnification is 665 × 665 *µm*^2^, two images are captured to visualize the whole channel width (1 mm) at each point. The fluorescence intensity profiles of each image are extracted with ImageJ after subtracting black noise and correcting for the inhomogeneous illumination profile. At each position, fluorescence intensity profiles are then stitched, normalized to the total fluorescence intensity of the image, and plotted with custom Python scripts.

The center of mass is computed from the fluorescence intensity profiles via Eq. 2 at each position and averaged over the four furthest positions from the inlet, to limit the effect of wash-out close to the inlet. The center of mass is then fitted with a linear function ⟨*Y*_*NM*_ ⟩(*t*) = *a* · *t* + *b* for *ϕ*_*M*_ *<* 0.17% or a saturating exponential 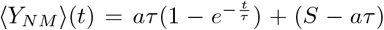 otherwise. The drift of the particles is the initial slope 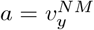.

### Measurements of bacteria drift

To measure the drift of both motile and non-motile particles, videos are acquired in the relevant fluorescence microscopy channel at different distances from the bottom surface and at least 100 *µ*m away from the lateral walls. To prevent the accumulation of non-motile cells on the sides, the videos are captured 30 seconds after the actuation of the flow, and then the flow is reversed for 30 seconds. The videos are then analyzed with the *φ*DM image velocimetry algorithm to extract the population-averaged drift in the field of view [75]. For measurements with density-matching media near the top of the microfluidic channel, we used particle tracking instead of *φ*DM image velocimetry to minimize the influence of particles below the focal plane, which can appear as background drift in the field of view and reduce the precision of velocimetry measurements.

### Swimming trajectories analysis with particle tracking

The trajectories of the rheotactic swimmers were measured on binary mixtures of a non-fluorescent nonmotile strain at 1.7% volume fraction and a non-tumbling motile strain at different volume fractions (0.51%, 0.17%, 0.051%, and 0.017%). Since particle tracking works best at low cell density, we mixed non-labeled and fluorescent motile cells, keeping the labeled volume fraction *ϕ*_*M*,labeled_ = 3.4 · 10^−3^ % constant. Using fluorescence microscopy, we captured videos of swimming cells away from the lateral walls, 2.6 *µ*m from the bottom surface for different imposed shear rates. Using a custom-made particle tracking ImageJ plugin [93], we analyzed the trajectories of the swimming bacteria and extracted the instantaneous orientation of the cells as *φ* = arctan(*v*_*y*_*/*(*v*_*x*_ − *v*_*P*_ (*z*_0_))), with *v*_*x*_ and *v*_*y*_ the instantaneous cell velocities in the x and y directions. We account for the flow advection by subtracting from *v*_*x*_ the Poiseuille flow velocity at *z*_0_ = 2.6 *µ*m, *v*_*P*_ (*z*_0_) = *v*_*x*,max_(2 ™ *z*_0_*/h*)*z*_0_*/h*, where *v*_*x*,max_ is the maximal advection speed measured at mid-channel height *z* = *h* = *H/*2 in the given condition.

### Unidimensional drift-diffusion simulations

In Fig. 1d, we employed a finite difference method to solve the advection-diffusion equation describing the system Eq. (1). We discretized the advection-diffusion equation with the Euler integration method as

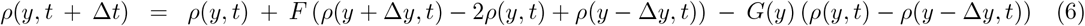

where

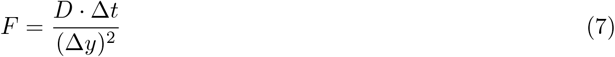

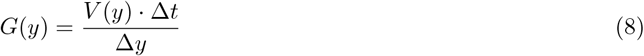

We set the initial density as *ρ*(*y, t* = 0) = 1, i.e. homogeneous across the spatial domain. We simulated the system for a total time of T = 7200 s, with a diffusion constant of D = 4 *µm*^2^*/s* and a total length of L = 1000 *µm*. The spatial grid spacing is Δ*y* = 5 *µm* and the integration time step Δ*t* = 1s. The velocity *V* (*y*) takes everywhere a constant value *v*_*y*_, i.e. that measured from the center of mass displacement for the given experiment, except in the first and last 50 *µ*m of the spatial domain, where it rises linearly from 0 to *v*_*y*_ to emulate the effect of lateral surfaces on the swimmer-induced flow. A fully constant velocity gave very similar results.

### Analytical model for backflow generation

The swimmers are modeled as a pusher dipole of point forces, one representing the body and the other the flagellum. The magnitude of both point forces is *f*_0_*/*6*πη* = *a*^*S*^*v*_0_, with *η* the fluid viscosity, *a*^*S*^ = 0.7*µ*m the hydrodynamic radius of the cell body, and *v*_0_ = 30*µm/s* the swimming speed. At low reynolds numbers, the fluid flow **v**^**i**←−**j**^ at position **r**_*i*_ generated by a point force **f**_*j*_ positioned at **r**_*j*_ is given by :

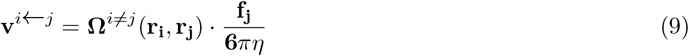

where **Ω**^*i*≠*j*^ is the normalized hydrodynamic-interaction tensor (the normalized Green function) of the incompressible Stokes equation:

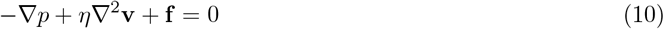

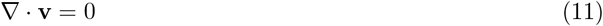

We model a section of the channel away from the lateral walls of the channel as a rectangular box of width and length W and height H = 2h, with open boundary conditions in the x and y directions and no-slip boundary conditions at the top and bottom walls. We take advantage of the 2D decomposition in Fourier space of the flow field between two parallel plates and its corresponding Oseen tensor 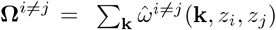 developed in [81, 82]. Under this partial Fourier decomposition, the incompressible Stokes equation reads:

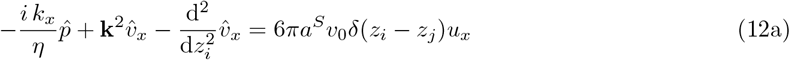

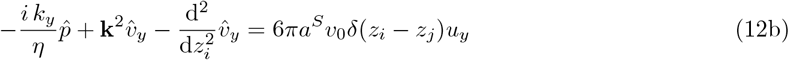

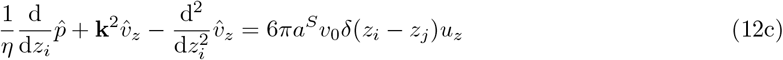

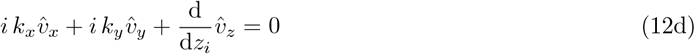

where **k** = (*k*_*x*_, *k*_*y*_) is the two-dimensional wave vector, *z*_*i*_ is the height of observation and *z*_*j*_ that of the point force, with *z*_*i*_, *z*_*j*_ ∈ [−*h, h*]. The Fourier transformed fluid velocity is 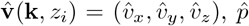 is the Fourier transformed pressure, the point force vector has an orientation **u** = (*u*_*x*_, *u*_*y*_, *u*_*z*_) and its norm is *f*_0_*/η* = 6*πa*^*S*^*v*_0_. Since we are interested in the average flow generated by the point forces, we consider the **k** = 0 component, 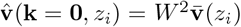, where 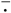 denotes the average over the *x* and *y* dimensions of space within the box.

The incompressibility Eq. 12d combined with the no-flow boundary conditions in *z*_*i*_ = ±*h* immediately give 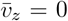. Hence, no average flow takes place in the vertical direction far from the lateral walls, and the vertical components of the force vectors get absorbed in average pressure gradients via

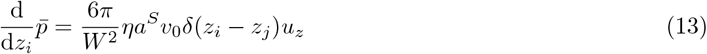

The horizontal components *α* = *x, y* of the average flow are then given by:

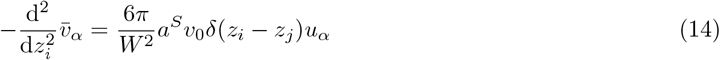

Following [81, 82], it is convenient to solve this equation by considering separately the cases *z*_*i*_ *> z*_*j*_ and *z*_*i*_ *< z*_*j*_ with their respective boundary conditions in *z*_*i*_ = *h* and *z*_*i*_ = −*h*, and then to stitch the two solutions by enforcing continuity in *z*_*i*_ = *z*_*j*_. This simple procedure yields:

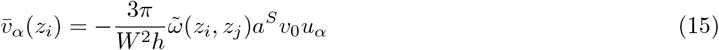

with

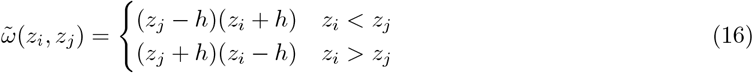

which can be written more synthetically as:

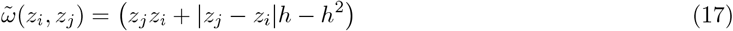

Note that in [81, 82], the authors impose a no-net-flow condition (∭ **v**d^3^**r**_*i*_ = 0) to the system by imposing a constant pressure gradient across the box, which adds a term to the expression of 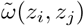. We do not impose this condition since we are precisely interested in the net flow induced by the swimmers. Moreover, because of its symmetry, this additional term has no net contribution to the problem at hand.

We consider a population of *N* swimmers, with swimming direction

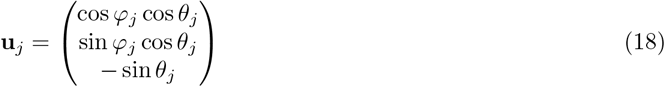

modeled as dipoles of point forces, where the cell body is located in *z*_*B,j*_ = *z*_*j*_ and produces the reduced force +*a*^*S*^*v*_0_**u**_*j*_, while the flagellum is located in *z*_*F,j*_ = *z*_*j*_ + *l*_*d*_ sin *θ*_*j*_ and produces the reduced force −*a*^*S*^*v*_0_**u**_*j*_. Because Stokes flows are additive, the mean backflow 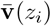 induced by this population reads:

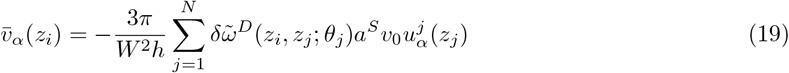

where the normalized dipolar Oseen tensor reads:

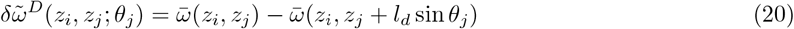

which, except in the small region *z*_*i*_ ∈ [*z*_*j*_, *z*_*j*_ + *l*_*d*_ sin *θ*_*j*_], reads:

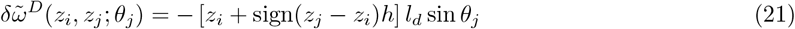

We now assume that the swimmers are distributed at the top and bottom surfaces in equal numbers, at heights *z*_*j*_ = *s*_*j*_(−*h* + *δz*_*j*_) with *δz*_*j*_ a small distance (typically one cell size) and *s*_*j*_ = 1 at the bottom, -1 at the top. Their orientation is given by the polar angle *φ*_*j*_ = *s*_*j*_*φ*(*j*), with *φ*(*j*) being drawn from the distributions of Fig. 4b, and a tilt *θ*_*j*_ = *s*_*j*_*θ*. The tilt angle towards the surface *θ* is taken constant here for simplicity, but the analysis is easily extended for a distribution of tilt angles. The multiplication by *s*_*j*_ accounts for the mirroring at the top of the channel of both the tilt and the distribution of *φ*, due to the change in vorticity direction for the latter. For *z*_*i*_ farther than max(*δz*_*j*_) + *l*_*d*_ sin *θ* from the walls, we can write, omitting the terms that cancel out when averaging over the motile population because they are of opposite sign when *z*_*j*_ is at the top or the bottom surface:

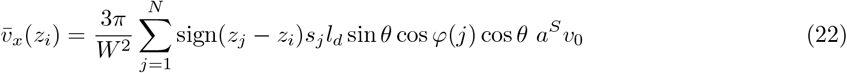

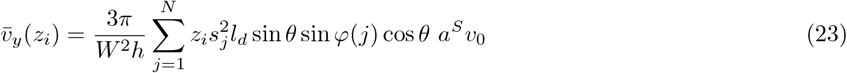

Introducing the motile cell number density *n*_*M*_ = *N/HW* ^2^ = *N/*2*hW* ^2^, and the average motile cell rheotactic drifts 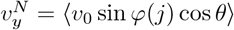, due to chirality induced rheotaxis, and 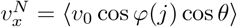 due to upstream surface rheotaxis, we obtain the advective backflows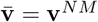:

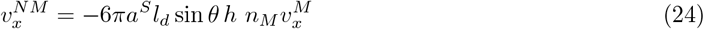

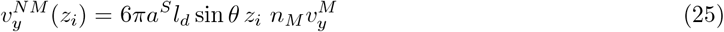

Finally, focussing on the chirality induced effect 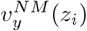, we compute the population averaged drift of non-motile cells in Fig. 4d,e as

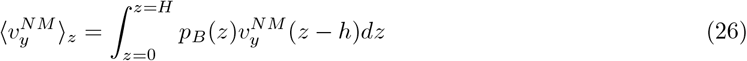

where *z* = *z*_*i*_ + *h*, and *p*_*B*_(*z*) is the vertical distribution of non-motile cells, which is skewed by sedimentation and is given by the Boltzmann distribution:

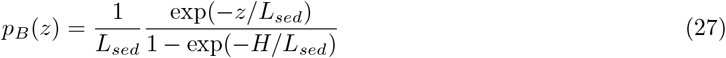

where *L*_*sed*_ = 8 *µ*m is the sedimentation length of non-motile particles for [14]. Integrating Eq. 26 with Eqs. 25 and 27, and thus neglecting the drop in velocity *v*^*NM*^ (*z*_*i*_) very close to the wall, we obtain the mean advection speed

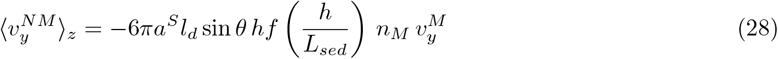

with

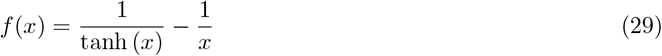

increasing from 0 to 1 as *h/L*_*sed*_ increases from 0 (density matched case) to infinity.

## Data availability

All data analysed during this study are included in this article and its supplementary information files.

## Code availability

Matlab code for the drift-diffusion simulations will be deposited on Gitlab https://gitlab.gwdg.de/remy.colin/uponpublication.

## Acknowledgments

We acknowledge support from the Deutsche Forschungsgemeinschaft, grant CO1813/2-1. We thank V. Sourjik for helpful discussion, Prof. Dr. K. Volz for access to the photolithography facility, G. Malengo for technical microscopy support, H. Henseling for building a custom microscopy stage and S. Espada Burriel for sharing strains.

## Author Contributions Statement

G.D.D. and R.C. designed the study and the experiments. G.D.D. performed the experiments. G.D.D. and R.C. analyzed the data and wrote the manuscript.

