## Supplementary information for "Active segregation in binary mixtures under flow"

February 20, 2025

### Supplementary note 1: Estimate of Darwin drift

If the motile cells were oriented parallel to the surface ($\theta = 0$), the average value of the instantaneous flow field induced by the dipole of force would be zero. Far from the swimmers, this would result in no net average transport of tracers, but for tracers that interact at close range with a swimmer, entrainment should occur as the speed of displacement of the tracer becomes non-negligible compared to the speed of the swimmer [1]. A body of finite size indeed drags fluid along as it moves. This is quantified by the so-called Darwin drift $V_D$, which is the net volume of fluid advected across the $x = 0$ plane by the body as it goes from $x = -\infty$ to $x = +\infty$ [2]. The concept was first introduced for (spherical) objects being advected by an external force (typically gravity), where $V_D > 0$ - i.e. fluid is advected forward. It was later extended to force-free microswimmers, where $V_D$ may become negative [1]. Since fluid entrainment results from short-range interactions between the swimmer and the advected fluid elements, quadrupolar terms in the flow field, which come from the different distributions of forces on the flagella and the body and from the finite size of the body, must also be considered [1, 3]. For such force-free swimmers, far from surfaces, the drift volume $V_D$ depends on quadrupolar terms in the flow field and on the wake-corrected volume of the swimmers $V_*$. Numerical simulations of bacteria similar to *E. coli* found $V_D \simeq -6(a^S)^3 = -1.5V_M$ for cells far from surfaces [1]. Close to a surface, fluid transport by a swimmer parallel to the surface is evidently affected by the latter, but the order of magnitude of the fluid displacement is not changed, although the sign might be, since quadrupolar contributions that entrain the fluid forward have a higher relative weight [3]. Finally, we can estimate the Darwin drift advection speed induced by a population of swimmers moving on average in the $y$ direction as $v_y^{NM} = n_M V_D v_y^M$ [4]. Therefore, a drift by this mechanism would be at most $|v_y^{NM,\text{Darwin}}| \sim 2\Phi_M v_y^M$, which is an order of magnitude lower than the drift we found $|v_y^{NM}| \simeq 20\Phi_M v_y^M$ [Fig. 2e]. This effect is thus very unlikely to be the main cause of the drift we observe.

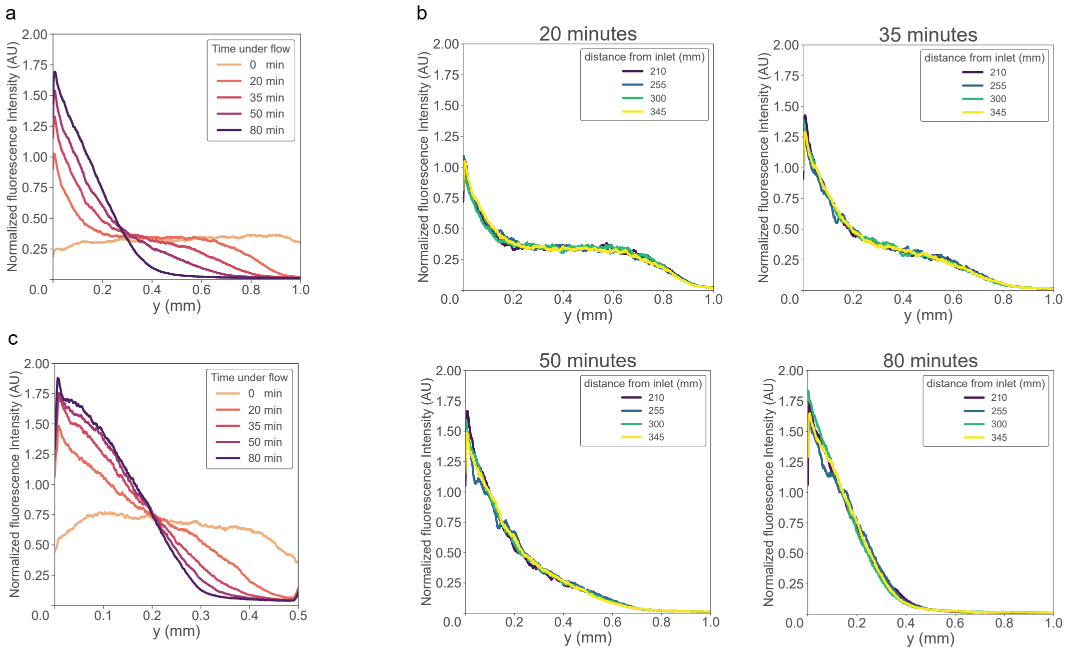


**Supplementary Fig. 1: Fluorescence intensity profiles at high non-motile cell concentration.** **a-c** Profiles of non-motile cell density distribution across the width of the channel, measured at 10.5 $\mu$m from the bottom surface from fluorescence images in experiments with high non-motile cell density ($\phi_M$ = 0.17%, $\phi_{NM}$ = 1.7% and shear $\dot{\gamma}$ = 5.6 $s^{-1}$) that are representative of $n$ = 3 biological replicates. **a**,**c** the profiles, which are shown for several time points in a channel of width W = 1 mm (**a**) or W = 0.5 mm (**c**), are averaged over the four last recording positions in the channel. **b** The profiles for the individual recording positions in the microfluidic channel of the same experiment as (**a**) are shown for different time points. The accumulation on the left side is independent of the position in the microfluidic chip and the width of the channel.

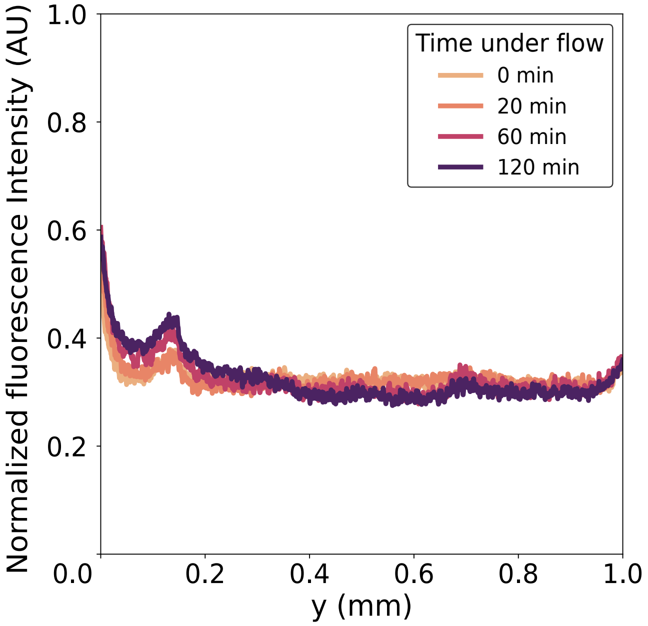


**Supplementary Fig. 2: The distribution of motile cells remains homogeneous in time under shear flow.** Distribution of motile cells across the width of the channel as it evolves over two hours, for an experiment with $\phi_M = 0.17\%$, $\phi_{NM} = 1.7\%$, shear $\dot{\gamma} = 5.6\ s^{-1}$, that is representative of the $n = 3$ biological replicates. The small increase in intensity at the left side of the channel (0 mm) is attributed to bleedthrough fluorescence from accumulating non-motile particles. The profile is the average of the density profiles over the last 4 positions in the chip.

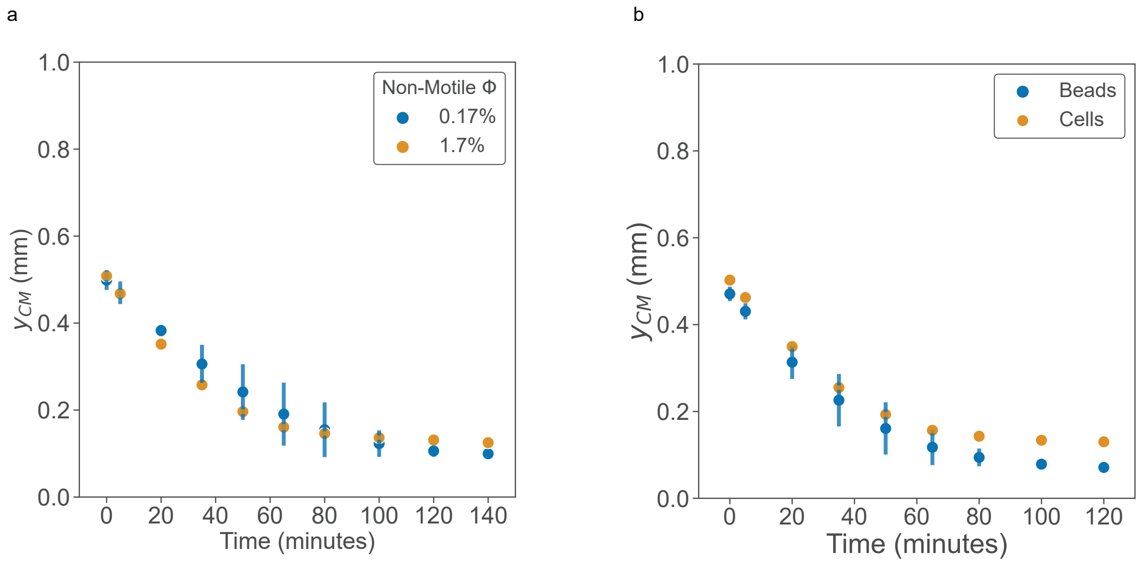


**Supplementary Fig. 3:** **a** Position of the center of mass of non-motile cells as a function of time since flow actuation for a high and a low concentration of non-motile cells ($\phi_M = 0.17\%$, shear $\dot{\gamma} = 5.6\ s^{-1}$). Error bars represent the standard deviation (SD) over $n = 3$ ($\phi_{NM} = 1.7\%$) or 2 ($\phi_{NM} = 0.17\%$) biological replicates. **b** Position of the center of mass of fluorescent passive colloidal beads (diameter $1.33\,\mu$m) as a function of time compared to that of non-motile cells. Experimental parameters are identical except for non-motile concentration ($\phi_{NM-Beads} = 0.34\%$, $\phi_{NM} = 1.7\%$, $\phi_M = 0.17\%$, shear $\dot{\gamma} = 5.6\ s^{-1}$). For the beads, error bars represent the standard deviation (SD) over $n = 2$ biological replicates.

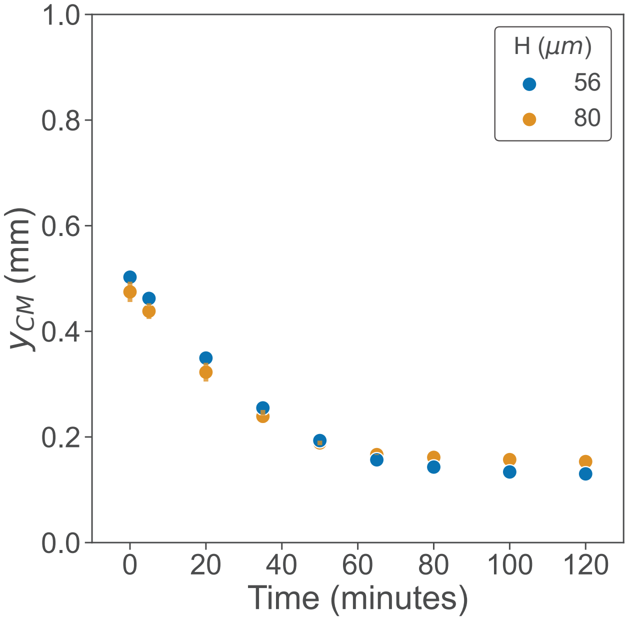


**Supplementary Fig. 4: Non-motile drift is independent of channel height at a fixed shear rate**. Position of the center of mass of non-motile cells over time in a channel with higher thickness H = 80 $\mu$m ($\phi_M$ = 0.17%, $\phi_{NM}$ = 1.02%, shear $\dot{\gamma}$ = 5.6 $s^{-1}$) compared to the standard channel with thickness H = 56 $\mu$m ($\phi_M$ = 0.17%, $\phi_{NM}$ = 1.7%, shear $\dot{\gamma}$ = 5.6 $s^{-1}$). Error bars represent the standard deviation (SD) over $n$= 3 (H = 56 $\mu$m) or 2 (H = 80 $\mu$m) biological replicates.

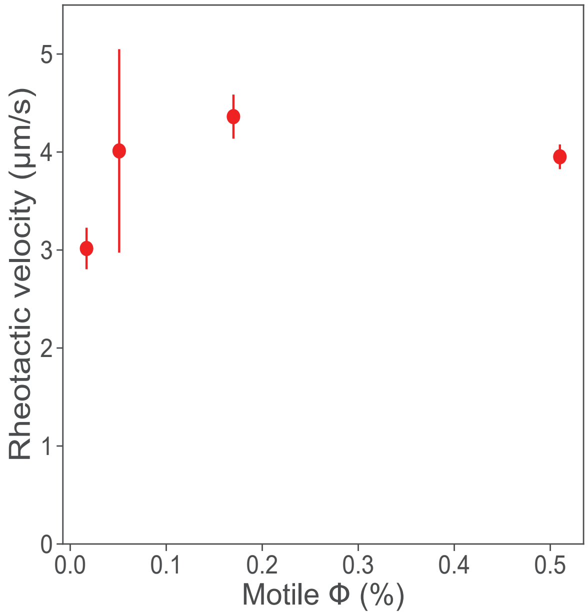


**Supplementary Fig. 5: Rheotactic drift of motile cells as a function of motile volume fraction.** For all the experiments, the volume fraction of labeled motile cells is kept constant at $3.4 \cdot 10^{-4}$ %, the rest being unlabeled, to facilitate tracking, the non-motile volume fraction is $\phi_{NM} = 0.17\%$ (unlabeled to prevent any bleedthrough), and flow shear is set at $\dot{\gamma} = 5.6\ s^{-1}$. Error bars represent the standard deviation (SD) over $n$= 3 biological replicates.

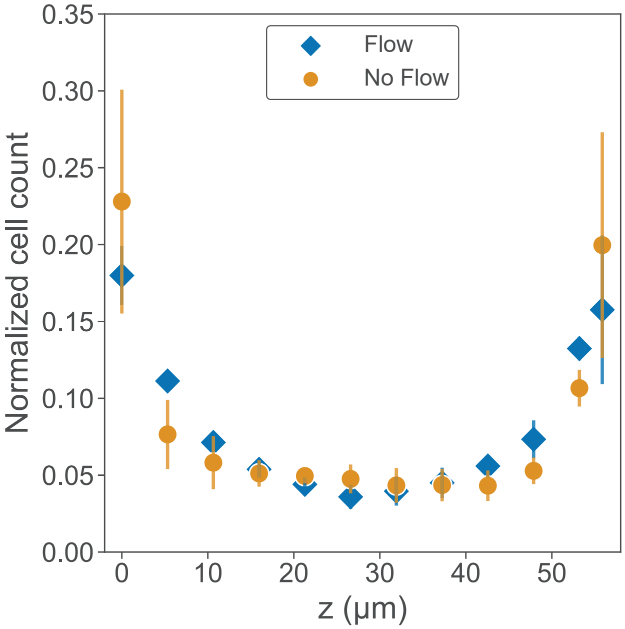


**Supplementary Fig. 6: Motile cells distribute predominantly near the top and bottom surfaces both in the absence and presence of flow.** Cell counts are computed from particle tracking within fluorescence images captured at each z-position in the absence of sedimentation. The cell count at each z-position is normalized to the total cell number across the channel for the respective biological replicate. For both experiments, the mixture is composed of $\phi_M = 0.17\%$, $\phi_{NM} = 1.7\%$ unlabeled cells, and $\phi_M = 0.0034\%$ labeled cells. The shear at the bottom is set at $\dot{\gamma} = 5.6$ s$^{-1}$. Error bars represent the standard deviation (SD) over $n$= 3 biological replicates.

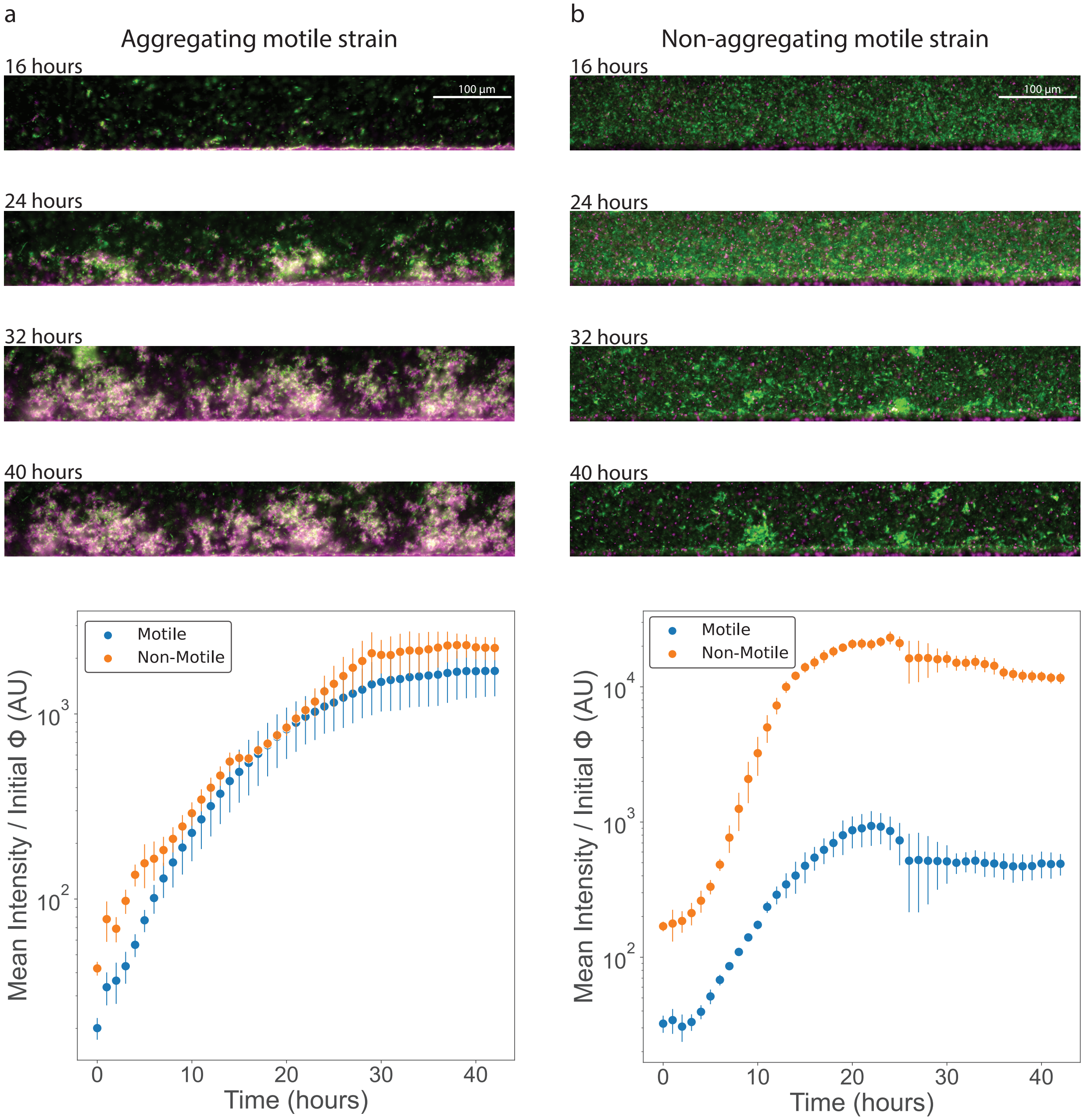


**Supplementary Fig. 7: Aggregating motile bacteria integrate more efficiently into asymmetric biofilms formed by non-motile bacteria than non-aggregating bacteria.** **a**,**b** Images (top) and quantification (bottom) of the bacterial biofilm at the left side of the channel as a function of time for a binary mixture of biofilm-forming non-motile bacteria (green) and motile bacteria (magenta), which are either able (**a**, wild-type) or unable (**b**, Δag43) to aggregate. Over time, aggregating motile cells are progressively integrated into the biofilm (**a**), whereas non-aggregating motile cells form an even mat on the bottom surface of the channel (**b**). **a**,**b** The mean fluorescence intensity of motile and non-motile cells was measured near the left side of the channel (667 $\mu$m x 150 $\mu$m) and normalized by the initial volume fraction of the respective strain, $\phi_M(0) = 0.17\%$ and $\phi_{NM}(0) = 0.051\%$. Applied shear is $\dot{\gamma} = 5.6$ s$^{-1}$. Error bars are standard deviation (SD) over $n = 2$ biological replicates.

| Name | Relevant genotype | Source |
|---|---|---|
| **Strains** | | |
| W3110 RH | wild-type | [5] |
| RC111 ($\Delta$*fliC*) | W3110 RH $\Delta$*fliC* $\Delta$*flu* | [6] |
| RC86 ($\Delta$*cheY*) | W3110 RH $\Delta$*cheY* $\Delta$*flu* | [6] |
| RC101 (WT) | W3110 RH $\Delta$*flu* pOB2; AmpR | [6] |
| RC125 ($\Delta$*cheY*) | W3110 RH $\Delta$*cheY* $\Delta$*flu* pOB2; AmpR | [6] |
| RC113 ($\Delta$*fliC*) | W3110 RH $\Delta$*fliC* $\Delta$*flu* pOB35; AmpR | [6] |
| RC114 ($\Delta$*fliC*) | W3110 RH $\Delta$*fliC* $\Delta$*flu* pOB2; AmpR | [6] |
| SEB21 ($\Delta$*fliC*) | W3110 RH $\Delta$*fliC* pOB35; AmpR | [6] |
| RC144 (WT) | W3110 RH pOB2; AmpR | [6] |
| **Plasmids** | | |
| pTrc99a | Expression vector; pBR ori, pTrc promoter, IPTG inducible; AmpR | [7] |
| pOB2 | *mCherry* in pTrc99A, IPTG inducible; AmpR | [6] |
| pOB35 | *mNeonGreen* in pTrc99A, IPTG inducible; AmpR | [6] |

**Table 1:** Strains that were used in this study. AmpR: ampicillin resistant
